# Nanopore Long-Read RNAseq Reveals Widespread Transcriptional Variation Among the Surface Receptors of Individual B cells

**DOI:** 10.1101/126847

**Authors:** Ashley Byrne, Anna E. Beaudin, Hugh E. Olsen, Miten Jain, Charles Cole, Theron Palmer, Rebecca M. DuBois, E. Camilla Forsberg, Mark Akeson, Christopher Vollmers

**Author notes:** Current Address: Department of Molecular and Cell Biology, School of Natural Sciences, University of California-Merced, Merced, CA, USA. Correspondence should be addressed to: C.V.

## Abstract

Understanding gene regulation and function requires a genome-wide method capable of capturing both gene expression levels and isoform diversity at the single cell level. Short-read RNAseq, while the current standard for gene expression quantification, is limited in its ability to resolve complex isoforms because it fails to sequence full-length cDNA copies of RNA molecules. Here, we investigated whether RNAseq using the long-read single-molecule Oxford Nanopore MinION sequencing technology (ONT RNAseq) would be able to identify and quantify complex isoforms without sacrificing accurate gene expression quantification. After successfully benchmarking our experimental and computational approaches on a mixture of synthetic transcripts, we analyzed individual murine B1a cells using a new cellular indexing strategy. Using the *Mandalorion* analysis pipeline we developed, we identified thousands of unannotated transcription start and end sites, as well as hundreds of alternative splicing events in these B1a cells. We also identified hundreds of genes expressed across B1a cells that displayed multiple complex isoforms, including several B cell specific surface receptors and the antibody heavy chain (IGH) locus. Our results show that not only can we identify complex isoforms, but also quantify their expression, at the single cell level.

## Introduction

Over the last decade, RNAseq has vastly increased our knowledge of eukaryotic gene expression and the unique transcript isoform signatures that differentiate developmental stages, organs, and single cells ^1–4^. Proteins that arise from transcript isoforms of a single gene can vary in their biological properties including stability, intracellular localization, enzymatic activity, and post-translational modifications ^5^. Transcript isoforms are the product of alternative <u>t</u>ranscription <u>s</u>tart <u>s</u>ites (TSSs), <u>t</u>ranscription <u>e</u>nd <u>s</u>ites (TESs), and alternative splicing events that include alternative splice sites, intron retention, and exon skipping ^6^. It has been predicted that a large fraction of human genes are alternatively spliced ^7,8^. Although alternative splicing enables increased transcriptome diversity, aberrations in splicing have been implicated in several human diseases, including cancer. Indeed, 15% of point mutations have been predicted to cause splice defects, resulting in human genetic disorders ^9^ and somatic mutations within splicing factors are associated with 12 different cancer types ^10^.

Consequently, it is important to determine the true transcriptional diversity of cells. This requires that gene expression is analyzed not only at the gene-level but also at the isoform-level. However, current short-read RNAseq methods are inherently limited in their ability to identify complex transcript isoforms, as they cannot sequence full-length transcripts. Instead, transcripts are fragmented for sequencing, resulting in short individual reads that fail to span the entirety of the transcript. Computational tools can be used to assemble full-length transcripts from these reads, but different assembly algorithms can result in conflicting outcomes and varying overall assembly quality ^11^.

To offset this limitation of short-read RNAseq, studies have successfully used both single-molecule long-read PacBio and synthetic long-read MOLECULO methodologies ^12–15^ to sequence full-length cDNA. However, PacBio technology has a bias toward shorter fragments necessitating the separation of cDNA by length before library preparation, which complicates sample preparation and quantification ^16^. Furthermore, MOLECULO depends on the assembly of short Illumina reads that suffer from the biases inherent in Illumina data and relies on the separation of individual transcript molecules into distinct wells. This complicates quantification as well as the analysis of highly abundant or similar isoforms. Recently, the Oxford Nanopore Technologies (ONT) MinION has been used to analyze full-length cDNA samples derived from both defined synthetic RNA molecules as well as RNA from tissue culture cells ^17^.

With the exception of a single study using single cell RNA-seq to focus its analysis on a single gene locus using PacBio technology ^18^, these long-read technologies have been used exclusively to evaluate transcriptome diversity across bulk cell populations. However, recent studies have highlighted that cells found within seemingly homogeneous populations can differ in gene expression ^19,20,21^. Understanding heterogeneity within cell populations has shown promise across multiple disciplines including developmental biology, neurobiology, cancer and immunology. Single cell approaches can help illuminate biological questions regarding cell function, development and dysfunction. Knowing the exact state of the cell can help determine its fate or reflect changes with response to stimuli or drug treatment, as well as its ability to neutralize a pathogen, respectively. Cell-to-cell heterogeneity ^3^ makes immune cells a fascinating target for in-depth analysis of transcriptional diversity. Current approaches that measure RNA transcripts within single cells rely on short-read RNA-seq, single molecule RNA-fluorescence in-situ Hybridization (SM-RNA FISH), or single-cell RT-qPCR ^22,23,24,25^. These current methods can either be applied to a few genes or are under the same constraints of short-read RNA-seq, which we described earlier. Ultimately, these approaches are unable to identify and quantify complex isoforms on a transcriptome-wide level.

To make it possible to identify and quantify complex isoforms on a transcriptome-wide single cell level, we have developed a nanopore sequencing approach for the analysis of full-length cDNA in single cells. The Oxford Nanopore Technologies (ONT) MinION sequencer is a portable device that is based on single molecule sequencing technology that provides reads of unprecedented length by performing voltage driven molecule translocations through small nanosensors ^26^. Although the MinION platform has been most useful for interrogating viral and bacterial genomes, recently it has been applied for analyzing cDNA in both targeted as well as genome-wide approaches ^17,27–30^. Taking advantage of its unprecedented read length, we wanted to interrogate single-cell transcriptomes of mouse B1a cells by sequencing full-length cDNA molecules using the ONT MinION sequencer.

We implemented an integrated informatics pipeline (*Mandalorion*) for gene-level and transcript isoform-level expression quantification to overcome the sequencing accuracy limitations of the ONT MinION. To identify transcript isoforms, *Mandalorion* predicted transcription start and end sites, as well as splice sites and their alternative usage. After benchmarking the ONT RNAseq approach on a complex mixture of synthetic transcripts, we sequenced seven individual mouse B1a cells and showed that we could accurately quantify gene expression and identify and quantify novel isoforms at the single-cell level. Our analysis identified differential usage of complex isoforms in over a hundred genes including several surface molecules like CD19, CD20, and IGH, the very receptors defining B cell identity.

## Results

### Generating and Sequencing Single-Cell RNAseq Libraries

We first investigated the ability of the ONT MinION platform to interrogate transcriptomes at the single-cell level. To test this, we used our ONT RNAseq approach to analyze seven individual mouse B1a cells ^31,32^ and compared it with the standard Illumina RNAseq approach. To this end, we FACS-sorted single B1a cells into individual wells containing lysis buffer and amplified cDNA from each individual cell using the Smartseq2 protocol with modifications (see Methods, Supplementary Table S1) ^33^. The cDNA generated by the Smartseq2 protocol was split and processed in-parallel using the Illumina and ONT library preparation protocols. Sequencing the fragmented cDNA of the seven cells on the HiSeq2500, we generated between 73,086-351,876 150bp Illumina reads per cell. Sequencing the full-length cDNA of the first three cells on individual ONT MinION flow cells using the R7.3 chemistry generated between 17,749-52,696 ONT 2D reads per cell (Supplementary Table S2). Taking advantage of the improved MinION throughput using the R9.4 chemistry, we multiplexed the full-length cDNA of the other four cells on a single MinION flow cell and generated between 57,874-128,726 ONT 2D reads per cell. To enable this multiplexing we introduced custom 60 nucleotide cellular indexes during PCR amplification (see Methods, Supplementary Table S1, Fig. 1a).

**Figure 1:**
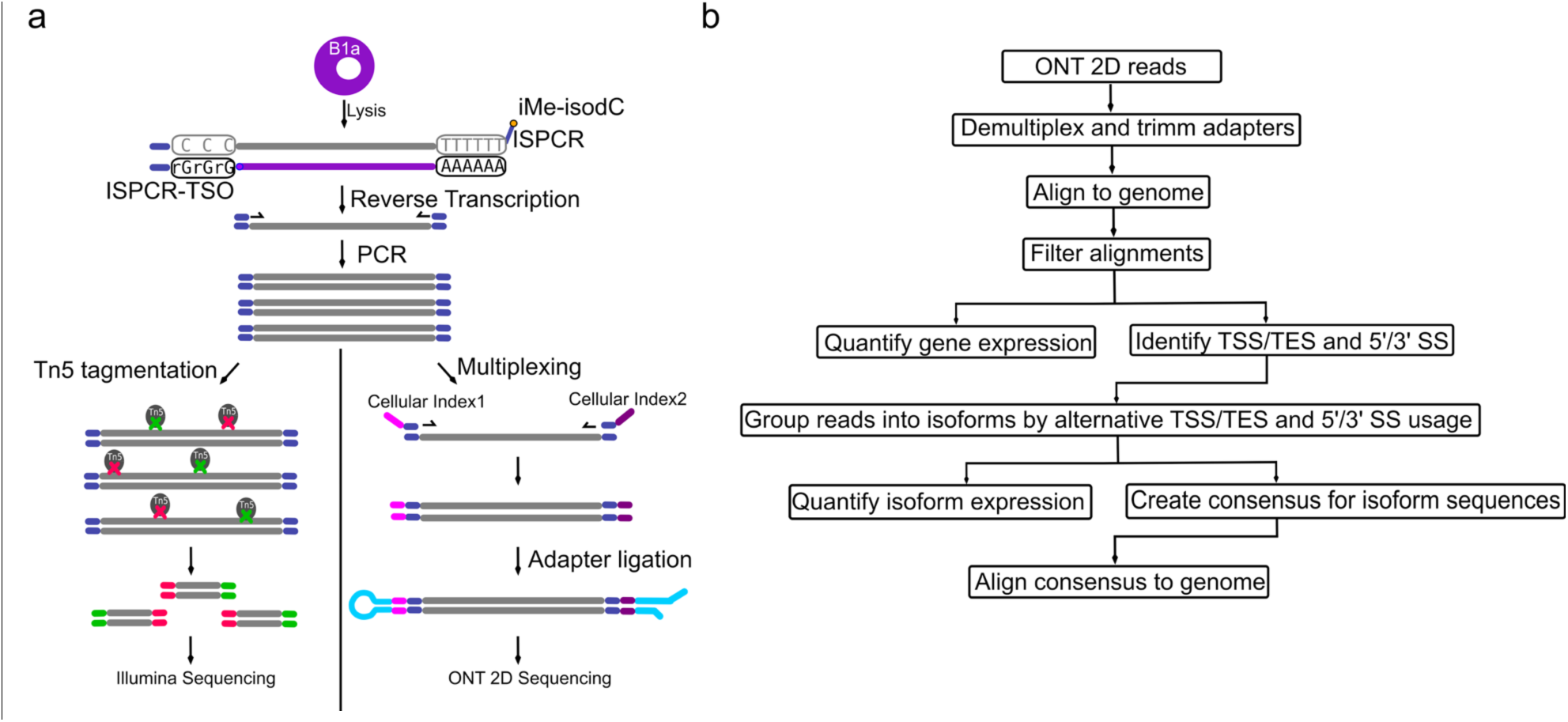
Experimental Design and Analysis Pipeline. a) Schematic of experimental design. FACS-sorted single B1a cells were lysed. PolyA-RNA was then reverse transcribed and PCR amplified using template switching. Full-length cDNA was split into two reactions. Half of the reaction was tagmented by Tn5 and seguenced using a Illumina HiSeg2500 seguencer. The other half of the reaction was ligated to ONT adapters and seguenced on an ONT MinlON seguencer. b) Schematic of the Mandalorion pipeline used to analyze the ONT 2D read data.

### Comparison of Gene Expression Quantification

To assess whether ONT RNAseq is capable of quantifying gene expression, we compared RNAseq data produced with ONT and Illumina, the current benchmark for gene expression quantification. Because standard gene quantification tools (eg. STAR^34^, Cufflinks^35^) are not compatible with nanopore reads, we aligned the ONT 2D reads using BLAT^36^ and quantified gene expression using our own *Mandalorion* algorithm. *Mandalorion* determines how many reads overlap with the exons of a gene to produce a <u>R</u>eads <u>P</u>er <u>G</u>ene per <u>10K</u> reads (RPG10K) value. As ONT 2D reads are long enough to span the full-length of the transcripts, normalization for gene length was not performed (Fig. 1b). Comparing Illumina and ONT RNAseq gene expression quantification for the same cell showed high correlation (Pearson r ≥ 0.84-0.89 for R7.3 and 0.9-0.92 for R9.4), confirming that our ONT RNAseq approach recapitulates Illumina gene expression quantification (Fig. 2). Comparing Illumina and ONT RNAseq gene expression quantification across different cells showed low correlation with a Pearson r ≤ 0.45, suggesting that ONT RNAseq can identify cell-to-cell variability ^1,37^ (Fig. 2).

**Figure 2:**
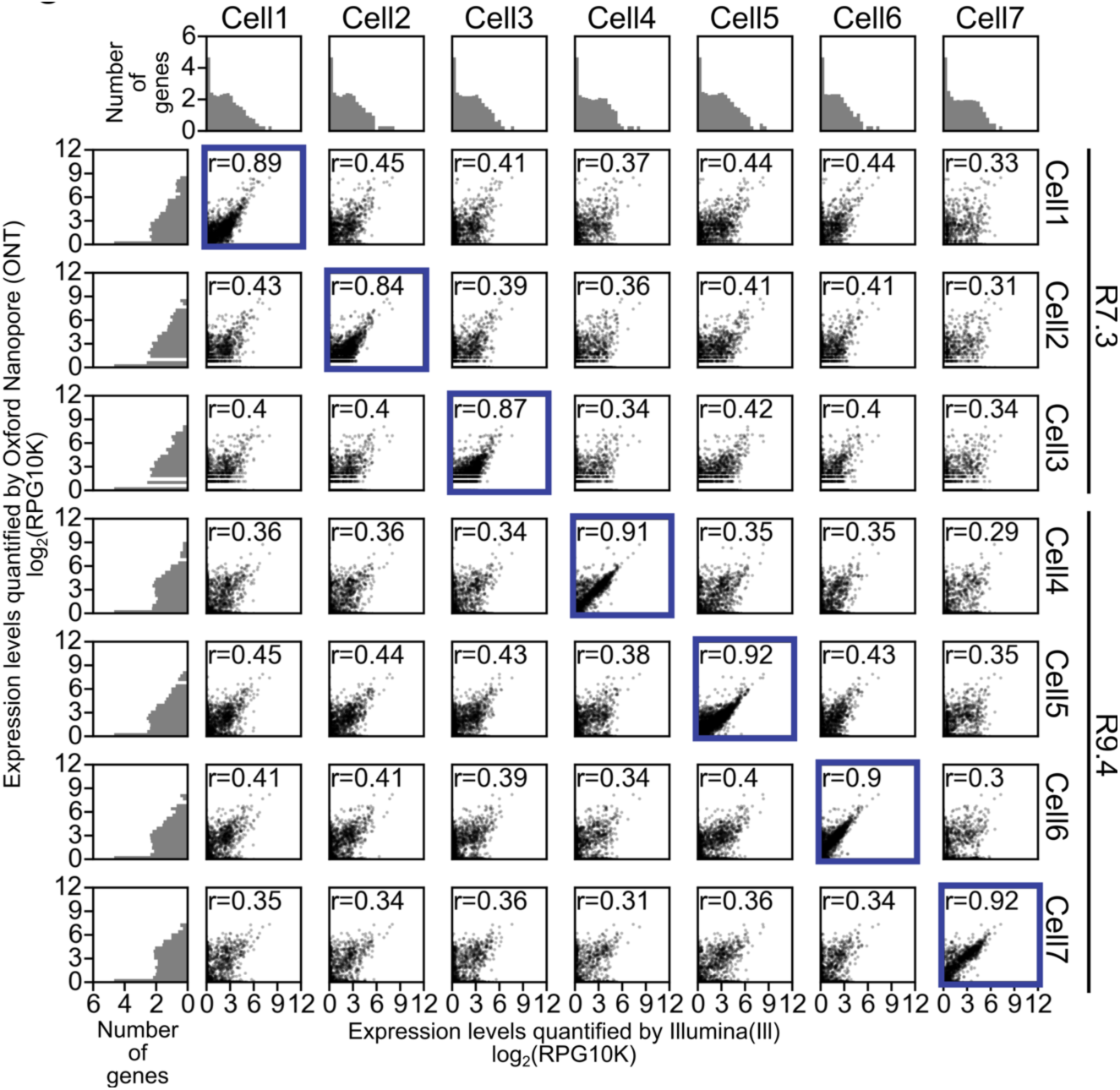
ONT RNAseq recapitulates Illumina RNAseq gene expression quantification. Scatter plot grid at the center of the figure shows gene expression levels for each gene as determined by Illumina RNAseq and ONT RNAseq for the indicated cells. Correlations of gene expression levels are given as <u>r</u>eads <u>p</u>er <u>g</u>ene per 10,000 reads (RPG10K) across 7 single cells. Pearson r is given for each cell per sequencing method combination with each point representing transcript expression level (x-axes =Illumina and y-axes=ONT). Same cell comparisons have a blue border. ONT sequencing chemistry is shown on the right. Histograms found on the left and top of the figure represent number of genes found binned by their expression levels.

These results show that even with the relative low number of reads produced, ONT RNAseq gene-expression quantification largely detects the same genes as Illumina RNAseq (Fig. 3a). Furthermore, subsampling ONT and Illumina raw reads showed that, for five of the seven cells analyzed, the detection of expressed genes had reached saturation (Supplementary Fig. 2). Unsurprisingly, genes that were detected by either ONT or Illumina RNAseq alone were expressed at lower levels, indicating that these genes were expressed at levels close to the detection limits of both technologies (Fig. 3b). We also observed that the genes detected by ONT RNAseq alone were comprised of smaller transcripts (Fig. 3c). Additionally, genes that were < 600 bp in length and were detected by both ONT and Illumina RNAseq had relatively lower expression levels in Illumina RNAseq data (Fig. 3d). While this is consistent with smaller transcripts being strongly selected against in the Tn5 based Illumina library prep, we couldn’t exclude that ONT RNAseq might have a bias towards shorter transcripts. To exclude this possibility, we chose to analyze a mix of synthetic transcripts.

**Figure 3:**
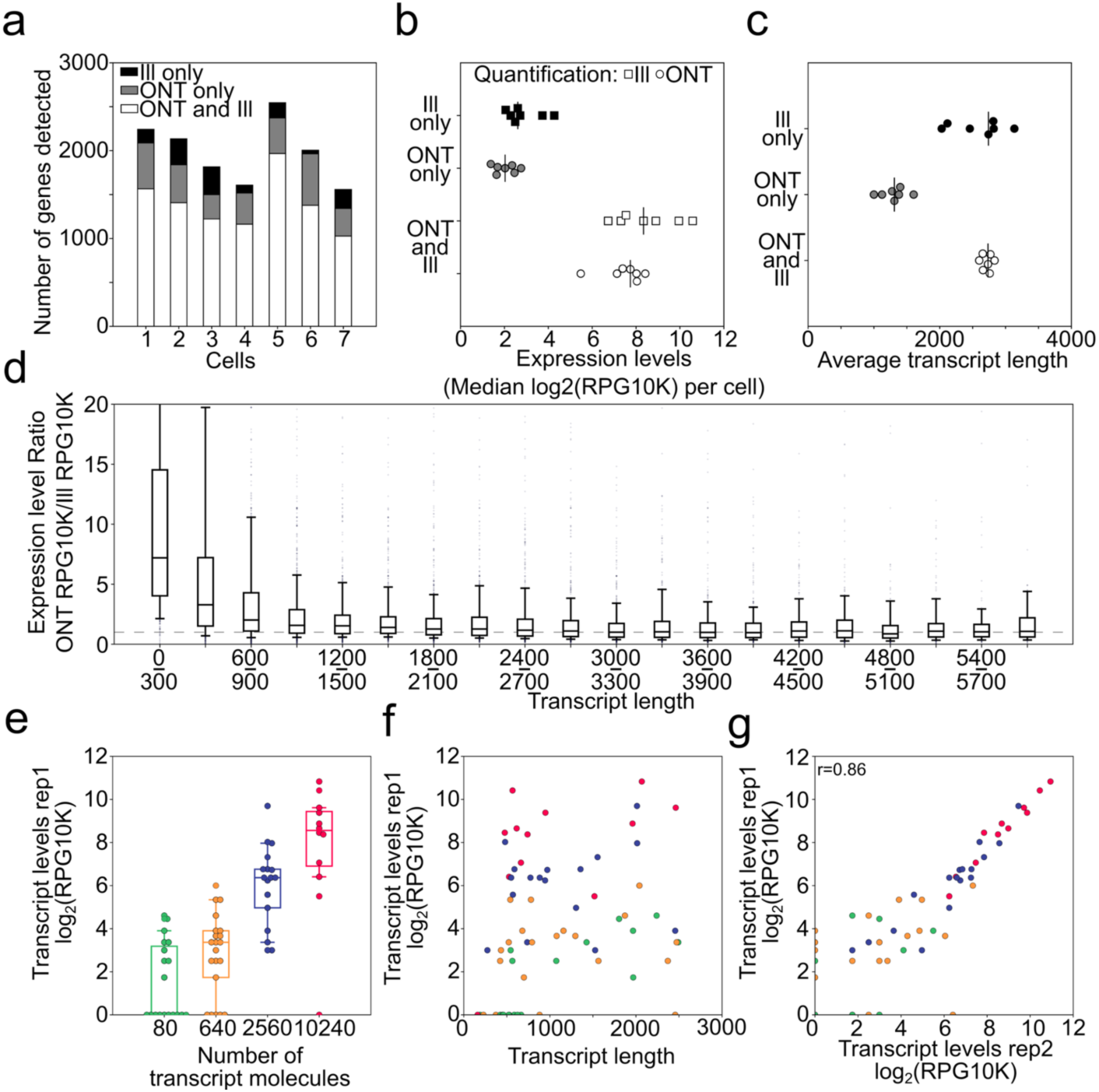
Quantifying gene and transcript expression with ONT RNAseq data. a) Stack barplots showing the number of genes detected by each cell corresponding to different sequencing technologies (III - Illumina, ONT - Oxford Nanopore). b) Median expression levels of genes detected by both or individual technologies. Two expression levels (III and ONT) are given for genes detected in both technologies. c) Gene length of genes detected by both or individual technologies. d) Ratio of gene expression levels for genes detected by both technologies. Ratios are binned according to gene length and shown as boxplots with whiskers indicating 10th and 90th percentiles. e) SIRV transcript levels of Replicate 1 (Rep1: 100fg SIRV pool E2) as measured with ONT RNAseq. Transcripts are binned by their starting molecule numbers. f) SIRV transcript levels of Replicate 1 are plotted against transcript length with colors corresponding to groups in e). g) Scatter plot showing correlation of SIRV transcript expression levels of Replicate 1 (Rep1: 100fg SIRV pool E2) and Replicate 2 (Rep2: 100fg SIRV pool E2), both measured by ONT RNAseq. r-value shown is Pearson-r.

### Analysis of synthetic transcript mixtures

To test whether transcript length had an effect on expression levels as measured by ONT RNAseq, we sequenced synthetic Spike-in RNA Variant Control Mixes (SIRVs, Lexogen) of known length, structure and sequence. SIRV transcripts provided in the E2 mix contained 69 transcripts ranging from 191-2528 nt. In the E2 mix 69 transcripts were present in four groups of varying concentrations containing 19, 21, 17 and 12 transcripts in each group, respectively. To test a wide range of possible transcript levels, we amplified (sub-) single cell amounts (i.e. 10fg and 100fg) of the Lexogen SIRV E2 mix in duplicate. This reflected a wide range of possible transcript levels with 8-10,240 molecules of individual SIRV transcripts present before the amplification step.

We quantified the 69 transcripts by aligning the resulting 5367-17915 2D ONT reads directly to the spliced SIRV transcriptome using BLAT and then counting and normalizing the matched ONT 2D reads for each transcript. As expected when amplifying (sub-) single cell amounts of RNA, we observed transcript drop-out in the lower concentration groups and found that transcript quantification showed variations within each concentration group (Fig. 3e, Supplementary Fig. 3a). Most importantly, however, quantification was not affected by transcript length, with the exception of transcripts shorter than 500 bp. These transcripts were either underrepresented or missed entirely (Fig. 3f). Generally, ONT RNAseq quantification agreed with the nominal concentration of the spike-in transcripts and, interestingly, the intra-group variations in transcript quantification were reproducible between replicates (Fig. 3g). This intragroup variation might be due to variation in initial transcript levels, systematic amplification bias, or data analysis bias. Overall, the observed underrepresentation of short transcripts in ONT RNAseq and the differences between Illumina and ONT RNAseq quantification are consistent with cDNA molecules below 500 bp in length being selected against during cDNA synthesis and again during the Illumina library preparation using the Tn5 method. Ultimately, analyzing these synthetic transcripts at different concentrations allowed us to exclude the possibility that ONT RNAseq favors shorter transcripts.

Next, we wanted to test whether, in addition to largely unbiased quantification of SIRV transcripts 500-2,500 bp in length, ONT RNAseq reads cover transcripts in their entirety which would make them uniquely suitable to identify and quantify complex isoforms.

### Using *Mandalorion* for genome annotation and isoform identification with SIRV ONT RNAseq data

The 69 synthetic SIRV transcripts are derived from 7 artificial gene loci that have been modeled after human genes with high isoform diversity, making them suitable for testing ONT RNAseq’s capability to capture isoform diversity in a genome annotation independent manner. To this end, we used *Mandalorion* to analyze ONT RNAseq 2D read data to annotate the SIRV gene loci, which in turn could be utilized to further identify and quantify SIRV isoforms. First, we used read alignments to annotate Transcription Start Sites (TSS) and Transcription End Sites (TES), as well as splice sites (SS) of SIRV transcripts in the SIRV gene loci. The annotation of TSS and TES was accomplished by end to end coverage of the entire RNA transcript by complete ONT 2D reads (i.e. reads for which both ISPCR adapters could be identified and trimmed, Supplementary Table 2) (Fig. 4a-c). Complete ONT 2D reads contained information regarding both TSS, TES in their read alignments.

**Figure 4:**
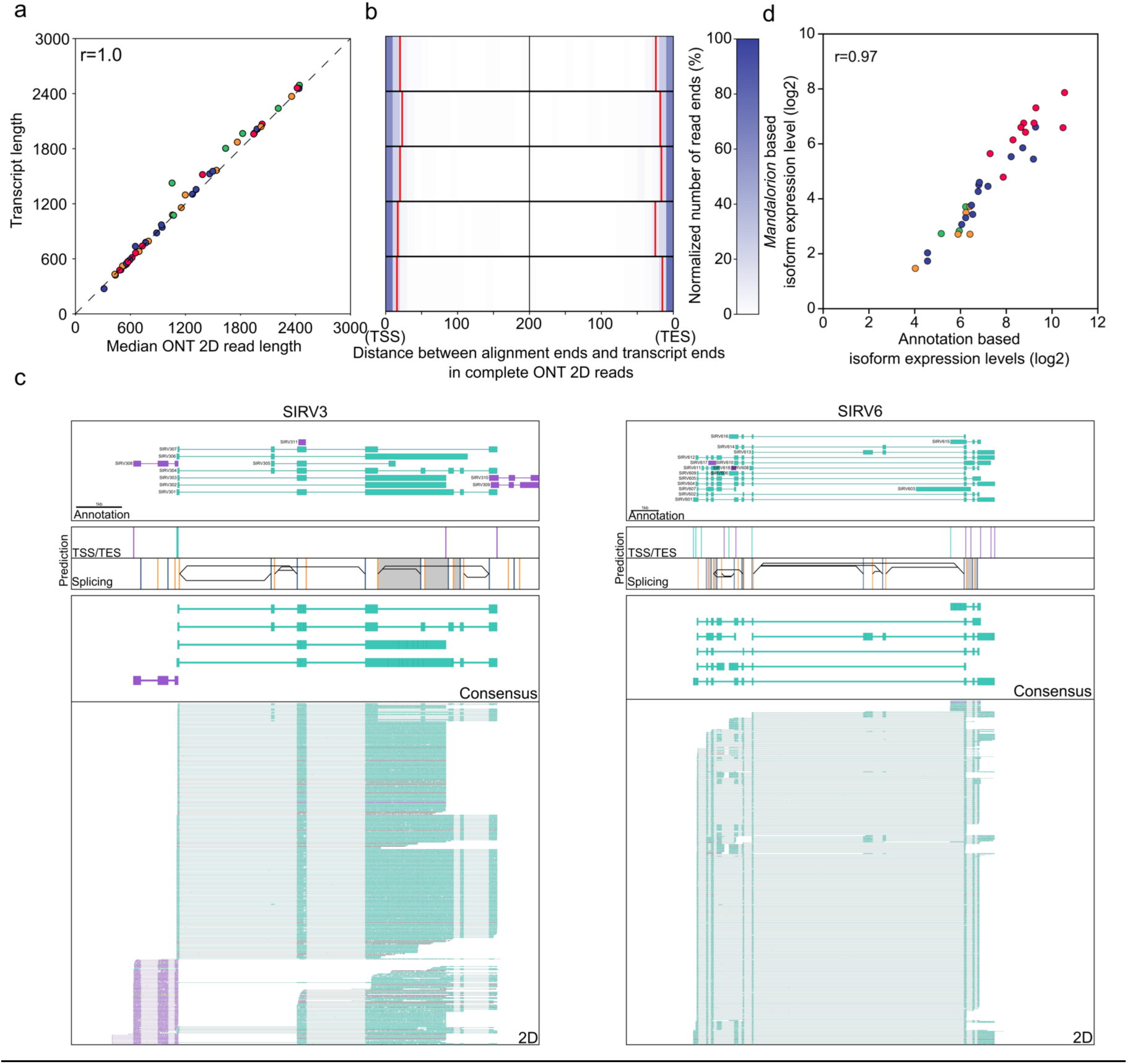
Identifying and quantifying complex isoforms in SIRV mixtures using ONT RNAseq data. a) Scatter plot shows correlation between ONT 2D reads and the SIRV transcripts they align to. Pearson r is shown. Coloring is as indicated in Figure 3e-g b) Distance between read alignment ends and transcript ends are shown as heatmap with the color indicating the normalized alignment numbers. 90% of read alignments terminated outside the red lines c-d) Genome Browser view of SIRV3(c) and SIRV6(d) gene loci. Top box contains transcript annotations, second and third box contain TSS (Teal) /TES (Purple) and splice sites (5’SS: yellow, 3’SS: blue) locations predicted from the read data, respectively. Black lines and grey areas in box 3 indicate alternative splicing and intron retention events predicted from the read data. Box 4 contains read alignments of isoform consensus reads. Box 5 contains ONT 2D read alignments. Direction of transcripts, isoform consensus, and ONT 2D reads are indicated by their color (Teal: 5’ to 3’, Purple: 3’ to 5’). e.) Scatter plot shows correlation between SIRV transcript quantification by aligning to annotated transcripts or by annotation-independent isoform grouping using Mandalorion. Pearson r is shown.

After combining and aligning ONT 2D reads of all replicates to the artificial SIRV genomic loci (Fig. 4c, Figure S4), we categorized 20 bp bins containing TSS, TESs and splice sites using the Madalorian pipeline (see Methods). To avoid the detection of spurious TSS and TES by prematurely terminated read alignments, we required TSS/TES to be at least 60 bp apart. In this manner, we detected 20 TSS and 24 TES that all directly overlapped with an actual TSS and TES and were within 60 bp of 38 (of 57) actual TSSs and 41 (of 59) actual TESs present in the SIRV transcript annotation. Furthermore, we detected 76 (of 89) 5’ splice sites and 73 (of 93) 3’ splice sites present in the SIRV genome annotation. By analyzing the actual splicing pattern of ONT 2D reads we detected 11 (of 12) alternative 3’ splice site combinations and 12 (of 14) alternative 5’ splice site combinations as well as 12 (of 12) intron retention events present in the SIRV transcripts.

Using *Mandalorion*, we then sorted ONT 2D reads into isoform groups based on their TSS/TES and alternative splice site usage. We generated consensus sequences of these groups using POA^38^ (Partial Ordered Alignment) and compared these consensus sequences to SIRV transcript sequences using BLAT. All of the 33 consensus sequences we generated matched a SIRV transcript with between 97.8% and 100% identity (BLAT identity score) and in all cases matched its directionality. Of the resulting 33 consensus sequences, 26 matched one of the 29 SIRV transcripts present in the two highest abundance groups (Fig. 4c, d, Supplementary Fig. 4). The other 7 consensus sequences matched one of the 40 SIRV transcripts in the two low abundance groups. While *Mandalorion* did not succeed in consistently identifying lower abundance isoforms, the consensus isoform sequences detected were very accurate. We also observed high correlation between the genome annotation-independent transcript isoform quantification by *Mandalorion* and quantification derived from directly aligning reads to the transcriptome (Fig. 4e) This means that in addition to identifying sequence, structure, and directionality of complex isoforms, *Mandalorion* can also accurately quantify them in a genome annotation independent manner. As a result, we were encouraged to apply this pipeline to our single cell data.

### Identification of Transcription Start and End Sites used in individual B1a cells

By analyzing the ONT 2D reads generated from the seven B1a cells using *Mandalorion*, we detected 4234 TSSs and 3883 TESs with only 2476 TSSs and 2448 TESs overlapping with the TSSs or TESs present in the Gencode annotation (vM10) ^39,40^ of the mouse genome (Fig. 5a). To determine whether the unannotated TSS and TES we detected were artifacts of our experimental and computational pipelines, we determined their Fantom5 ^41^ CAGE peak and polyA signal enrichment. Fantom5 CAGE peaks are derived from capturing and sequencing the 5’ end of transcripts and should therefore be enriched in TSSs. Indeed, we found that in contrast to TESs (49/3883 or 1.3%), a high percentage of both annotated (2356/2476 or 95%) and unannotated (1052/1799 or 58%) TSSs overlapped with high scoring Fantom5 CAGE peaks (Fig. 5b). Conversely, both annotated and unannotated TESs were highly enriched for polyA signals, while TSSs were not (Fig. 5c). When we assigned the detected TSSs and TESs to annotated genes, we found that most genes contained exactly one TSS and one TES, as expected. However, 696 genes contained more than one TSS or TES indicating the presence of more than one isoform (Fig. 5d). Overall, this suggested that *Mandalorion* successfully identified thousands of unannotated TSSs and TESs and hundreds of genes with differential TSS/TES usage by analyzing individual cells.

**Figure 5:**
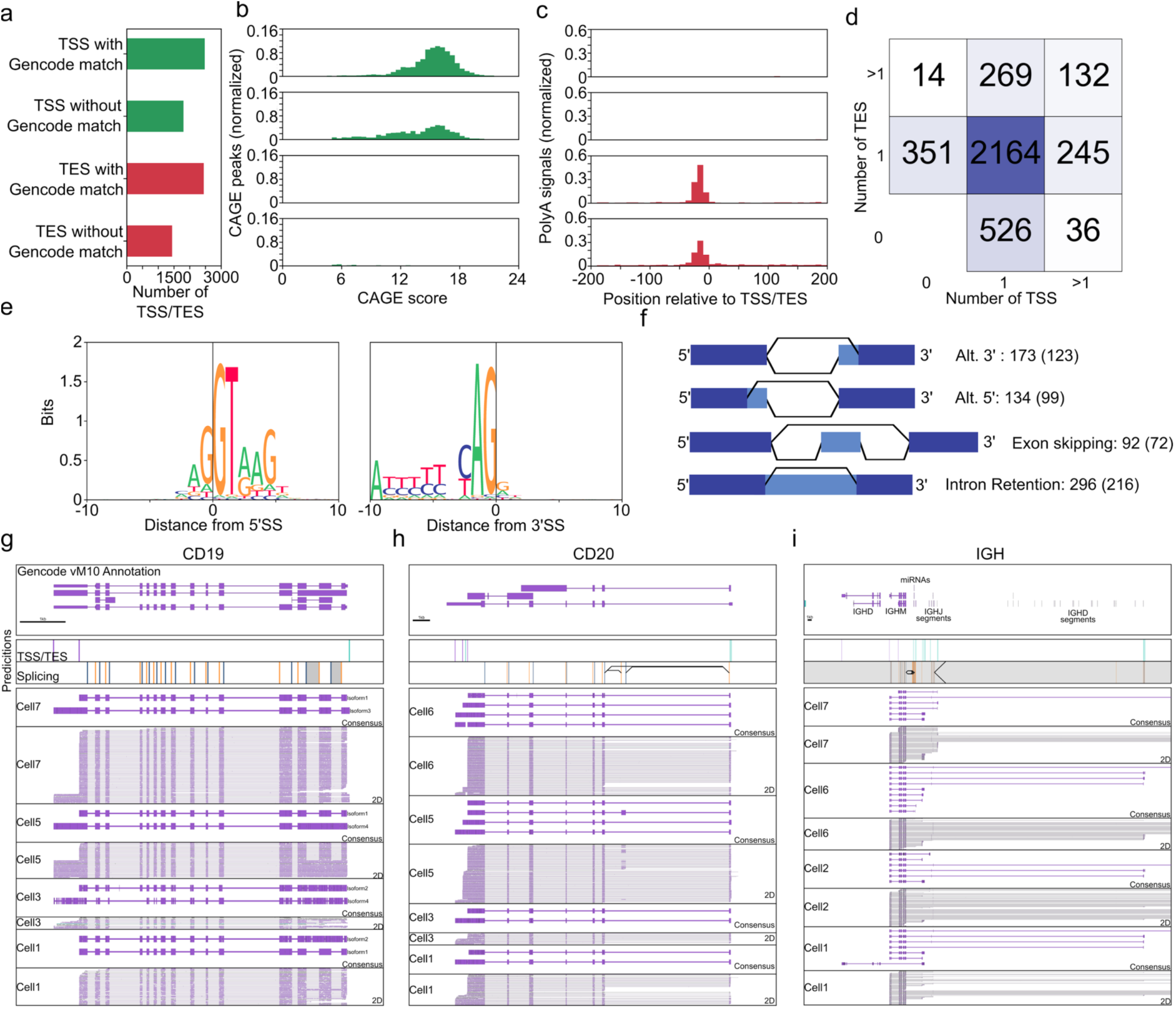
Mandalorion analysis of ONT RNAseq data identifies transcription start and end sites, splice sites, and isoforms in mouse B1a cells. a) TSSs and TESs predicted based on read data were separated into sites with or without GENCODE vM10 annotation matches. b-c) TSSs/TESs with or without GENCODE matches were tested for FANTOM5 CAGE area enrichment (b) and polyA signals (c). d) Overlap of TSSs and TESs with genes. Genes were sorted according to the number of TSSs and TESs they overlapped with. e) Predicted base composition at 5’ and 3’ SS based on read data is shown as sequences logos. f) Schematic for detection and corresponding number of detected alternative splice site combinations. g-i) Genome Browser view of CD19, CD20, and IGH gene loci as shown in Figure 4. ONT 2D reads and consensus sequence alignments are shown for the indicated cells. Splice sites for the highly repetitive IGH locus were not considered for isoform grouping due to the difficulty of aligning reads unambigiously.

### Identification of Alternative Splicing Events used in individual B1a cells

In addition to TSSs and TESs, *Mandalorion* identified a total of 24,887 5’ splice sites and 24,756 3’ splice sites. The vast majority of these splice sites were supported by the GENCODE annotation or splice junctions found in Illumina reads. Of the 24,887 5’SS and 24,756 3’SS we identified, 24,298 (97.6%) and 24,220 (97.8%) matched GENCODE annotation, respectively. Of the 589 5’SS and 536 3’SS that did not match GENCODE annotation, 250 (42.4%) and 216 (40.2%) were supported by splice junctions in Illumina reads, respectively. Even if all splice sites that were not supported by GENCODE annotation or Illumina reads were false, which is unlikely, the false discovery rate of our approach would only be 1.3% (659/49,643).

Furthermore, while *Mandalorion* defined our splice sites as 20 bp bins, we were relatively successful in defining the exact splice site as shown by the base context of the determined splice sites (Fig. 5e). By determining alternative splice sites, we found 296 intron retention events, 134 alternative 5’ splice sites and 173 alternative 3’ splice site combinations. The majority of these events were also observed in Illumina read data, which supported 216 (of 296) intron retention events, 99 (of 134) alternative 5’ splice sites, 123 (of 173) alternative 3’ splice sites and 72 (of 92) exon skipping events (Fig.5f). Alternative events not supported by Illumina read data had significantly lower ONT 2D read counts than those that were supported (Table S3), indicating they might be closer to the detection limits of both technologies.

### Identification of Complex Isoforms

Having established that ONT RNAseq can be used to identify isoform features like TSSs and TESs as well as alternative splicing events, we aimed to identify complex isoforms. We defined genes as expressing complex isoforms if they contained alternative TSS/TES as well as alternative splice sites. We identified 169 genes that expressed complex isoforms. By identifying and quantifying all isoforms we detected at these 169 genes, we found highly significant differential isoform usage between cells in 55 of the genes (Chi2-contigency test, alpha=0.001, holm-sidak multiple-testing correction). These genes with significant differential isoform usage included B cell specific surface receptors CD19 and CD20, the antibody heavy chain locus (IGH) (Fig. 5g,h,i), CD37 (Fig. 6), as well as CD2 and CD79b, and CD45 (Supplementary Fig. 5). We created consensus sequences of the isoforms at these gene loci in each B1a cell and found that across the individual B1a cells, isoforms derived from CD19 showed a combination of alternative TSSs and intron retention events. Isoforms derived from CD20, on the other hand, showed a combination of alternative TESs, as well as an exon skipping event including a previously unannotated exon. The IGH locus was even more complex, with canonical isoforms containing VDJ recombinations and the IGHM constant region exons. In addition, we observed isoforms containing the IGHM constant region exon but originating from 1.) abortive DJ recombinations 2.) I-exon 3.) miRNA loci in the IGHM Switch-region, and 4.) a J-segment. Finally, one isoform in cell 1 originated from the IGHM I-exon but contained the IGHD constant region exons. While IGH isoform diversity has been previously observed and has been known for a long time to be involved in class-switching ^42^, the ability of ONT RNAseq to sequence full-length cDNA at the single cell level truly highlights and confirms the exceptional transcriptional diversity of the IGH locus.

**Figure 6:**
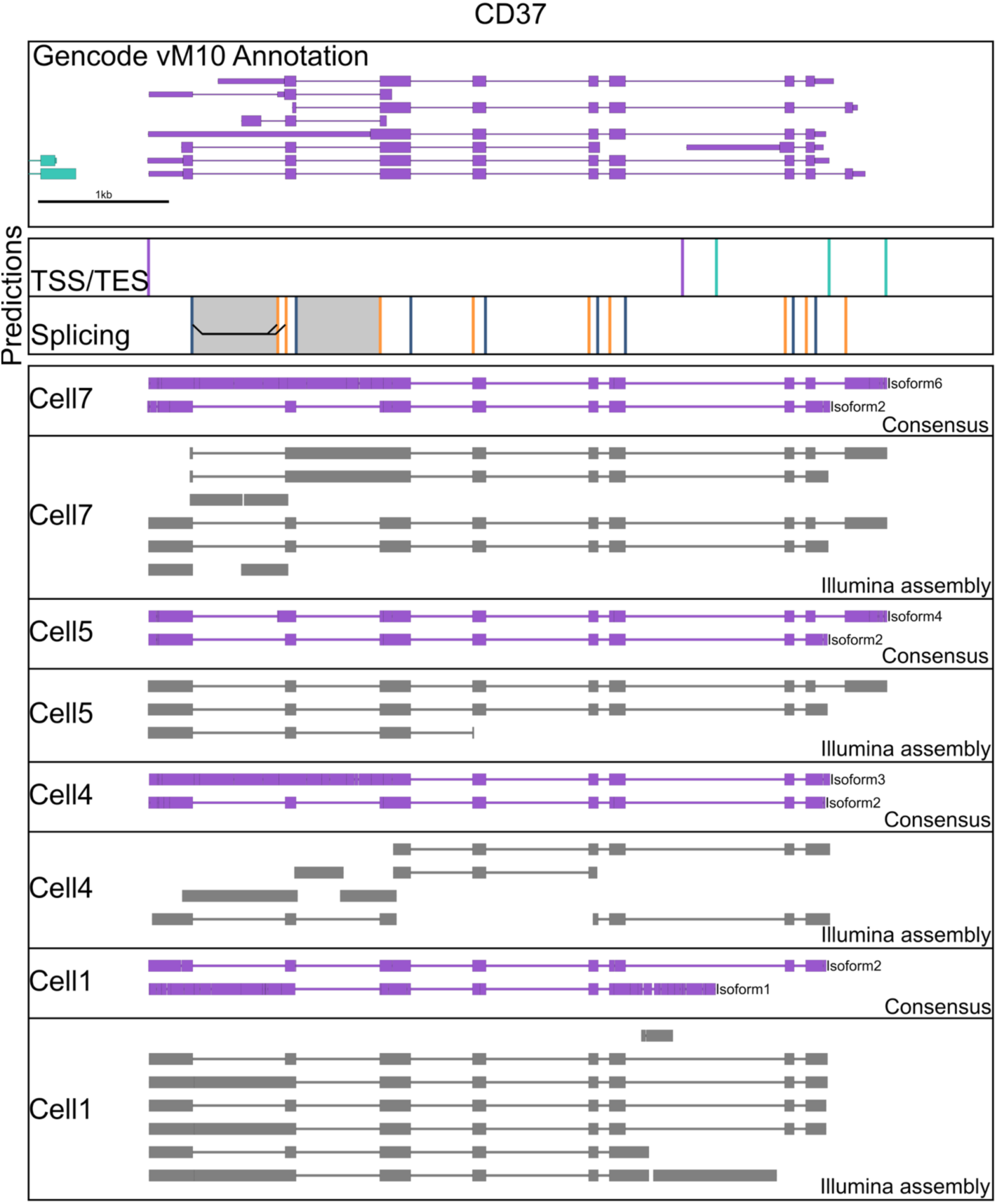
Uncovering isoform diversity in B cell surface receptors. Genome Browser view of the CD37 is shown as in Figure 4. In addition to isoform consensus derived from ONT 2D reads, contigs assembled from Illumina data using Trinity are shown in grey for the indicated cells.

The ability to sequence entire cDNA molecules from end to end presents an advantage over assembling transcript isoform using Illumina data. While assembling Illumina data using Trinity ^43^ is likely to succeed if a gene locus only expresses a single isoform, it appears to struggle with analyzing multiple isoforms of a gene locus that contain multiple distant alternative features. For example, ONT RNAseq identified several distinct isoforms of the CD37 gene across the individual cells analyzed. In most cases, when we assembled the Illumina data from individual cells, Trinity was either unable to form complete contigs or produced contigs that were shown by ONT RNAseq to be misassembled (Fig. 6). The CD37 gene and its isoforms therefore highlight the strength of the ONT RNAseq approach to identify the diversity of complex isoforms beyond what is possible with either bulk or short reads technologies.

## Discussion

The data we present here shows that RNAseq studies using the Oxford Nanopore Technologies MinION sequencer have the potential to redefine the level of information gathered by a single RNAseq experiment.

By benchmarking our experimental and computational pipelines on ONT MinION data derived from a mix of synthetic transcripts, we showed that our approach identifies the location of transcription start and end sites as well as splice sites in a genome. Furthermore, we have shown that these experimentally determined annotations can then be used by our *Mandalorion* pipeline to identify and quantify complex isoforms longer than ∼500 bp in an otherwise largely transcript length independent manner. It is likely that if we use less stringent size selection methodologies during library preparation, we could capture transcripts < 500 bp as well. Although we were only able to consistently identify the SIRV transcripts found among the high abundance groups, we expect that the less abundant transcripts could be identified using our pipeline by increasing the sequencing depth. Variation in the quantification of transcripts in the SIRV mix indicated that quantification might be improved by using <u>U</u>nique <u>M</u>olecular <u>I</u>dentifiers (UMI)^44^ during cDNA amplification. However, UMI length would have to be at least >30bp to be resolved unambiguously with the current error-rate of the ONT MinION. Introducing random nucleotides of this length during priming is likely to create short, unwanted PCR artifacts which would greatly increase the noise of the amplification reaction. Ultimately, until ONT sequencing accuracy improves, the Smart-seq approach employed in this study is currently the best choice for UMI free library generation, as it has been shown by a comparison study to generate the smallest amount of PCR duplicates and the highest transcriptome coverage when comparing low input methodologies ^45^.

By focusing on single cells transcriptomes, we demonstrated the capability of sequencing read output and accuracy of ONT MinION sequencer. We showed that ONT RNAseq can not only quantify known genes with a high correlation to Illumina RNAseq but, using the *Mandalorion* pipeline we developed, also annotate transcript features, thereby allowing us to identify and quantify complex, never before observed, isoforms. Using ONT RNAseq on only seven B1a cells, we identified thousands of unannotated transcription start and end sites which we then validated using FANTOM5 CAGE data and polyA signals, respectively. Furthermore, we identified 696 genes displaying alternative transcription start and end site usage, and 354 genes with alternative splicing events. Although not all alternative splicing events we detected were supported by single cell Illumina data, the events that weren’t supported were of significantly lower coverage, indicating they might be closer to the detection limits in either technology (Supplementary Table 3). Combined with the relatively low Illumina sequencing depth per cell in our study, this suggests that larger Illumina depth might aid in the validation of individual events in future studies.

In addition to the identification of individual alternative events, the read length of the ONT MinION sequencer paired with our *Mandalorion* analysis pipeline enabled us to identify 169 genes expressing complex isoforms containing both alternative TSS/TES and splice sites. Interestingly, among the genes expressing these complex isoforms were surface receptors, including the very surface receptors distinguishing B cells from other immune cells. For example, we found that CD19, CD20 (Ms4a1), IGH, CD45 (B220 or Ptprc), CD2, CD79b, and CD37 were expressed as multiple complex isoforms across the seven B1a cells. This indicates that the diversity of the surface receptors found on B-cells is not fully understood, which could have important implications on all facets of B cell biology. Our data suggest that we are currently only scratching the surface of the true transcriptional diversity of B1a cells. In the future, we aim to use the multiplexing strategy that we have developed to analyze hundreds of individual cells. This will make it possible to truly reconstitute the full transcriptome complexity of B1a and other cell types and will likely lead to discovery of additional subpopulations with distinct functional properties ^1^. While we currently estimate the cost per cell at ∼ $100-200, this is likely to decrease considering the rapidly increasing throughput of the ONT MinION and the soon-to-be-released ONT PromethION sequencer.

Nanopore sequencing is still rapidly maturing and we believe that advancements in sequencing chemistries, nanopore design and analysis algorithms will vastly improve the technology and address the shortcomings of low read numbers and high error rates in the near future. Lower error-rates will, for example, allow us to improve the *Mandalorion* pipeline further by enabling the base accurate identification of TSS/TES and splice sites, instead of identifying 20 bp bins for these features. Even with its current limitations, the data and analysis tools we present here demonstrate the potential of ONT RNAseq to revolutionize analysis of transcriptomes. Finally, while the ONT MinION has not quite caught up with the very capable PacBio Sequel long read sequencer, it is only a fraction of its price (∼$1,000 vs. $300,000). At this price, any molecular biology lab will be able to perform their own RNAseq experiments onsite, thereby increasing adoption of the single cell RNAseq approach and accelerating research.

## Methods

### FACS sorting of individual B cells

Mice were maintained in the UCSC vivarium according to IACUC-approved protocols. Single murine Ter119^−^CD3^−^CD4^−^CD8^−^Gr1^−^B220^+^ IgM^+^CD11b^−^CD5^+^ B1a cells were isolated from wild-type C57Bl/6 mice by lavage and incubated with fluorescently-labeled antibodies prior to sorting ^31^. The following antibodies were purchased from Biolegend to stain B-cells: Ter119, CD3 (145-2C11), CD4 (GK1.5), CD8a (53-6.7), B220 (RA3-6B2), Gr1 (RB6-8C5), IgM (RMM-1), CD5 (53-7.3), and CD11b (M1/70). Cells were analyzed and sorted using a FACS Aria II (BD), as described previously ^46,47,48^. Single cells were sorted into 96 well plates and directly placed into 4 ul of Lysis Buffer - 0.1% Triton X-100, 0.2 ul of SuperaseIn (Thermo), 1ul of oligodT primer (IDT), 1ul of dNTP (10mM each)(NEB) - and frozen at -80°C.

### Smartseq2 cDNA synthesis

Single cell lysate was reverse transcribed using Smartscribe Reverse Transcriptase (Clontech) in a 10 ul reaction including a Smartseq2 ^33^ TSO (Supplementary Table S1) according to manufacturer’s instructions at 42°C. The resulting cDNA was treated with 1 ul of 1:10 dilutions of RNAse A (Thermofisher) and Lambda Exonuclease (NEB) for 30 minutes at 37°C. A PCR amplification step using KAPA Hifi Readymix 2x (KAPA) step was performed incubating at 95°C for 3 mins, followed by 27 cycles of (98°C for 20 s, 67°C for 15 s, 72°C for 4 mins), with a final extension at 72°C for 5 mins.

### Illumina Sequencing

The resulting full-length cDNA PCR product was treated with Tn5 enzyme ^49^ which was loaded with Tn5ME-A/R and Tn5ME-B/R adapters (Supplementary Table S1). The Tn5 product was then nick-translated and amplified for 13 cycles (72°C for 6 mins, followed by 98°C for 30s and 13 cycles of (98°C for 10 s, 63°C for 30 s, 72°C for 2 mins), with a final extension at 72°C for 5min) with KAPA Hifi Polymerase (KAPA) and Nextera Index Primers (Supplementary Table S1). Libraries were then size selected using a E-gel 2% EX (Thermo-Fisher) to a size range of 400-1000 bp and sequenced on an Illumina HiSeq2500 2×150 run.

### Nanopore Sequencing

To achieve the 1ug of DNA needed for the Oxford Nanopore library prep, the full-length cDNA product was split into five aliquots and amplified for 13 cycles with KAPA Hifi Readymix 2X (KAPA) using the ISPCR or multiplex cellular index primers. The following reaction was incubated at 95°C for 3 mins, followed by 13 cycles of (98°C for 20 s, 67°C for 15 s, 72°C for 4 mins) with a final extension at 72°C for 5 mins. The single cDNA or multiplex product was further end-repaired and dA-tailed using NEBNext Ultra End Repair/dA tailing mix (NEB), and adapter ligated using the sequencing adapters provided by ONT (HP Adapter/Adapter Mix). Ligation reaction was performed using Blunt/TA ligase master mix (NEB). Reactions were then enriched using Dynabeads MyOne C1 Streptavidin (Life Technologies) to capture molecules that contain the HP Adapter. Enriched libraries were then mixed with Fuel mix and Running buffer provided by ONT. Single cell libraries were either sequenced solely on one (Cell1 and Cell2) or two (Cell3) separate MinION R7.3 flow cells and ran on the 48 hr 2D protocol. For our multiplexing strategy, single R9.4 flow cells were used (Pool1: Cells4-7, Pool2: Lexogen libraries) and ran on the 48 hr 2D protocol.

### Data Analysis

#### Illumina data

Illumina paired end 150 bp reads in fastq format were quality and adapter trimmed using trimmomatic (v0.33) ^50^. The trimmed reads were aligned using STAR (v2.4) ^34^ to the mouse genome (Sequence: GRCm38, Annotation: gencode vM10 ^40^). Illumina reads were assembled for each cell separately using the Trinity (v2.2.0)^43^ set of tools.

#### ONT data preprocessing

ONT reads were processed using the Metrichor cloud platform 2D workflow. For R7.3 runs, both reads that passed or failed Metrichor quality cutoffs were retained. For R9.4 runs, reads that failed Metrichor quality cutoffs were discarded as they also failed our alignment criteria. Fast5 files generated by Metrichor were converted into fastq and fasta formats using poretools (v0.5.1) ^51^.

#### ONT data analysis

##### Demultiplexing and adapter trimming (*Mandalorion*)

For demultiplexing, index-sequences were aligned to the reads using BLAT with parameters: - noHead -stepSize=1 -minScore=20 -minIdentity=20. Reads for which index-sequences could be identified were trimmed and assigned to the respective libraries. Next, for multiplexed and non-multiplexed reads alike, ISPCR adapter sequences were identified and trimmed using Levenshtein distances. Reads for which ISPCR adapters could be identified and trimmed were marked but all reads, trimmed or not, were aligned to the mouse genome (GRCm38) using BLAT(v35×1)^36^ with parameters: -stepSize=5 -repMatch=2253 -minScore=100 -minIdentity=50 - maxIntron=2000000. Alignments were filtered for a single alignment per read. This filtering process involved three steps: (i) the highest scoring alignment for each read is identified, alignment scores within 2% of each other were treated as ties, (ii) in case of ties the alignment with largest number of gaps is selected (this selects against alignment to unspliced pseudogenes) and (iii) if the best alignment of a read has a ratio of aligned bases to read bases ≤0.6 the read and its alignment are discarded.

##### Gene Expression (Mandalorion)

Gene Expression for ONT and Illumina RNAseq was analyzed using custom scripts. For each gene, the number of reads overlapping with its exons was counted, normalized to total number of aligned reads in a library and reported as <u>R</u>eads <u>P</u>er <u>G</u>ene per 10,000 reads (RPG10K). Genes were counted as expressed if they had a RPG10K value>0. RPG can be calculated as:

> *RPG10K = (total# of reads aligned to a gene’s exons ÷ total # of aligned reads in sample)×10,000*

##### Transcription start and end site detection (Mandalorion)

For the detection transcription start and end sites we limited our analysis to reads for which we detected and trimmed ISPCR adapter prior to read alignment. We then identified positions in the genome at which at least 2 alignments of these complete reads ended. We then further restricted our analysis by only considering positions with a median and 75th percentile of the number of clipped (unaligned) read bases between 6-15 and ≤ 20, respectively. This number of clipped (unaligned) bases corresponds to the length of bases contained in ISPCR TSO and oligodT primers that were not trimmed. We then placed a 20 bp bin around these positions to include the highest number of read alignment ends possible.

To filter false positive bins caused by incomplete read alignments in highly expressed genes bins were only considered as containing true TSS/TES if they met the following conditions: i) The total number of read alignment ends in the bins had to be ≥ 2% of the total number of reads in the next 50 read covered bases. ii) The candidate site had to be at least 60 read covered bases away from the next closest TSS/TES.

By only counting bases covered by read alignments we didn’t take non-covered introns into account which would skew our analysis. Next, in order to distinguish TSS and TES bins, we calculated median Levenshtein distances of the unaligned bases at all read alignment ends in a bin to nucleotides present in TSO (ATGG) or the OligodT (TTTT) primer. If the median Levenshtein of a bin to ATGG was ≤2 it was declared a TSS. If the median Levenshtein of a bin to TTTT was ≤2 it was declared a TES.

##### Transcription start and end site validation

To assess Fantom5^41^ CAGE scores we downloaded combined CAGE data (mm9.cage_peak_phase1and2combined_coord.bed.gz), converted the data to mm10 coordinates using https://genome.ucsc.edu/cgi-bin/hgLiftOver ^52^ and investigated direct overlap between TSS/TES and CAGE peaks. We considered TSS/TES and CAGE peaks to be overlapping if they were within 10 bp of each other. To assess polyA enrichment in TSS/TES, we extracted genomic sequences up- and downstream of these sites and looked for identical string matches to “AATAAA” and “ATTAAA”.

##### Splice Site detection(Mandalorion)

To identify 20 bp bins as splice sites only ONT 2D reads with a ratio of aligned bases/read bases of > 0.9 were analyzed. We then identified positions in the genome at which at least two read alignments of these reads opened or closed an alignment gap larger than 50 bp.

The 20 bp bins surrounding these positions were considered as containing a splice site if the following conditions are met: To filter false positive bins caused by spurious read alignment gaps in highly expressed genes bins were only considered as containing true splice sites if they met the following conditions: i) the number of of reads opening or closing an alignment gap in the bin was at least 2% of the total number of reads in the preceding (5’) or subsequent (3’) 40 read covered bases. ii) not closer than 30 bp to another splice site. The directionality of the splice site bin containing either 5’ or 3’ status was based on the direction of the majority of reads containing the splice site.

##### Alternative Splicing (Mandalorion)

To detect alternative splice sites, we counted how often 5’ and 3’ splice sites were spliced together in ONT 2D reads with aligned bases/read bases ratio of > 0.85. A 5’->3’ combination had to be present in at least 2 reads to be considered. We scored alternative splice site usage if the same 5’ splice site was spliced into two different 3’ splice sites or vice versa. To detect intron retentions, we identified areas between 5’ and 3’ splice sites that were covered to at least 70% by at least one ONT 2D read.

##### Isoform identification and quantification (*Mandalorion*)

We detected isoform by grouping reads according the TSS/TES and alternative splice sites they contained. ONT read alignment ends found within 60 bp of a TSS and a TES were sorted based on which alternative splice sites it contained. Isoforms that contained at least 1% of all reads at a gene locus were retained. All the reads in these retained isoform groups were used to create consensus reads using POA^38^. In short, fasta files containing all read sequence are passed to POA which generates a consensus of the reads by creating a multiple sequence alignment of the reads in the form of a partially ordered graph. The program then returns the most heavily-weighted path as the consensus of the reads. The consensus reads are then aligned to genome using BLAT parameters: -stepSize=5 -repMatch=2253 -minScore=10 -minIdentity=10. There was however, one exception regarding the highly complex variable regions derived from the IGH transcripts which were first aligned with IgBlast ^53^ and then with BLAT. IgBlast alignment coordinates were converted to genome coordinates and BLAT and IgBlast portions of the read alignments were merged.

##### Statistical test and multiple testing correction

We used the ‘chi2_contingency’ function in the scipy.stats ^54^ package to implement the Chi2 contingency test to detect differential expression of complex isoforms between cells. Multiple testing holm-sidak correction was performed with the ‘statsmodels.sandbox.stats.multicomp’ ^55^ package.

#### Data Visualization

All data analysis and visualization was performed in python ^56^ using the numpy/scipy/matplotlib ^54,57,58^ packages.

## Data Availability

Illumina and ONT sequencing reads were uploaded to the SRA under accession number SRP082530. All scripts, including the *Mandalorion* pipeline, are available upon request and will be available at https://vollmerslab.soe.ucsc.edu/

## Acknowledgments

Research reported in this publication was supported by NIH/NHGRI awards HG006321 (M.A.) and HG007827 (M.A.); by the NIH/NIDDK award R01DK100917 (E.C.F.); by Alex’s Lemonade Stand Foundation Innovation Award (E.C.F.); by the American Asthma Foundation (E.C.F.); by CIRM Training grant TG2-01157 to A.E.B.; by NHLBI K01HL130753-01A1 to A.E.B.; by CIRM Shared Stem Cell Facilities CL1-00506 and CIRM Major Facilities FA1-00617-1 awards to UCSC. E.C.F. is the recipient of a California Institute for Regenerative Medicine (CIRM) New Faculty Award RN1-00540 and an American Cancer Society Research Scholar Award RSG-13- 193-01-DDC. A.B. and C.C. are funded by the NHGRI/NIH training grant 1T32HG008345-01.

## Author contributions

C.V., M.A., and E.C.F. conceived of the research. C.V. directed the research. C.V., A.B., and C.C. analyzed the data. C.V. and A.B. wrote the paper. A.B., A.E.B., T.P., M.J. and H.O. designed and performed experiments. R.M.D. contributed materials. All authors edited the manuscript.

